# Investigation into *Nicotiana* systemin orthologs reveals lineage specific systemin insensitivity in LAB accession *N. benthamiana*

**DOI:** 10.64898/2026.11.13.766558

**Authors:** Euan W. McCubbin, Adam D. Steinbrenner

## Abstract

Systemin (SlSYS1) from *Solanum lycopersicum* was the first plant peptide hormone to be identified, and its receptor (SYR1), processing, and role in defense are all well characterized. However, studies of systemins outside of *Solanum* are extremely limited. Here, we identified four new bioactive systemins within the genus *Nicotiana*, three from *N. tabacum*, NtSYS4A/B/C, and one from *N. benthamiana*, NbSYS4. We showed that expressing the canonical systemin receptor SlSYR1 confers the ability to detect these novel systemins. In addition, we found that LAB accession *N. benthamiana* contains a premature stop codon in its SlSYR1 orthologs, providing a potential basis for its insensitivity to systemins. We showed that *N. tabacum* roots can respond to NbSYS4, and that expressing its SYR1 homolog NtSYR1A allows for the detection of systemins in *N. benthamiana*. Additionally, we found that NtSYS4A can downregulate NtSYR1A-dependent ROS, akin to antisystemin (SlAntiSYS) in tomato. Our findings challenge the consensus that systemins are limited to the *Solanoideae* clade and uncover a new immune signaling system within *Nicotiana*.

## Introduction

Phytocytokines are a diverse class of immunomodulatory peptides involved in activating defense responses to pathogens and herbivores (Hou et al. 2021). Phytocytokines are translated as longer precursor proteins that are proteolytically processed to release the mature, bioactive peptide. Some phytocytokines are highly conserved, such as ProPEPs and RALFs, which were first discovered in *Arabidopsis thaliana* and *Nicotiana benthamiana* respectively, but have been shown to mediate defense responses throughout flowering plants (Huffaker et al. 2013, Liu et al. 2024). However, many phytocytokines have only been shown in specific lineages, such as the *Zea mays–*specific ZIP1 (Ziemann et al. 2018), and GmPep890 and GmPep914, which are largely specific to *Glycine max* (Yamaguchi et al. 2011). The low degree of sequence conservation among phytocytokines, even within the same family (Rhodes et al. 2021), often obscures evolutionary origins.

Systemin (SlSYS1), an 18-amino acid phytocytokine initially isolated from *Solanum lycopersicum*, was the first endogenous signaling peptide to be discovered in plants (Pearce et al. 1991). Systemin is synthesized in plants as Prosystemin (PS), an intrinsically disordered 200 amino acid (aa) protein, lacking a signal peptide. Prosystemin enters the apoplast, likely via mechanical disruption of the plasma membrane, where it is cleaved via a phytaspase (Beloshistov et al. 2018), releasing the mature systemin peptide. Systemin is then detected by the cell surface receptors, systemin receptors 1 and 2 (SYR1/2), which are both leucine-rich repeat receptor-like kinases (LRR-RLKs). SYR1 binds to systemins with high affinity, triggering the recruitment of SERK3 coreceptors, leading to a strong immune output. At high concentrations of systemin the low-affinity, but more abundant receptor SYR2 outcompetes SYR1 for SERK3s, attenuating the immune response (Zhou et al. 2025).

In addition to the canonical systemin SlSYS1, multiple other homologs have been described. Within *S. lycopersicum*, a recent study found the immunogenic paralogs SlSYS2/3/4, the antagonistic paralog SlAntiSYS, and the physiologically uncharacterized paralog SlSYS5 (Wang et al. 2025). The *S. lycopersicum* prosystemin (SlPS) proteins vary substantially in length ranging from ∼50 to 200 aa. Differential signaling between SlSYS peptides has been shown to be mediated by their differential recruitment of SERK co-receptors, homologous to Arabidopsis BAK1. SlAntiSYS binding to SYR1 fails to recruit SERK3, and SlSYS1 recruits SERK3 but at a lower affinity than SlSYS2 does (Wang et al. 2025, Wang et al. 2026).

Investigations into PS and SYR homologs outside of tomato have been notably sparser. PS homologs have long been identified within *S. nigrum, S. tuberosum, Capsicum annuum* (Constabel et al. 1998) and *Physalis ixocarpa* (Hernández et al. 2008), but apart from a recent investigation into *S. tuberosum*’s StSYS1 and its role in viral defense (Skripnikov et al. 2026) no systemin outside of *S. lycopersicum* has been characterized. Notably the study which first described SYR1/2 also identified members of *Nicotiana* as having “SYR-like” orthologs (Wang et al. 2018). However, there has been no prior research into these putative receptors beyond genomic identification.

Here we describe four new *Nicotiana* PS homologs, three within *N. tabacum* we term NtPS4A/B/C and one within *N. benthamiana* we term NbPS4. NtSYS4B/C and NbSYS4 are bioactive against *N. benthamiana* expressing SlSYR1. NbPS4 undergoes wounding-induced protease-dependent processing in a manner similar to but likely distinct from SlPS1-4 in tomato. We also found that *N. tabacum*, and wild accessions of *N. benthamiana* have full length SYR orthologs, but not the widely used LAB accession. In line with previous studies, none of the systemins described caused an immune response in LAB *N. benthamiana*; however, wild-type *N. tabacum* roots and seedlings responded to some but not all systemins. Expressing the *N. tabacum* SYR ortholog NtSYR1A in LAB *N. benthamiana* confers the ability to respond to all four known bioactive SlSYS peptides and all four described *Nicotiana* SYS peptides. Interestingly, pretreating NtSYR1A-expressing *N. benthamiana* with NtSYS4A attenuates its response to NbSYS4 in a similar manner to SlAntiSYS with SlSYR1-expressing *N. benthamiana*, providing a potential mechanism for systemin signaling regulation in *Nicotiana*. Our findings reveal new aspects of the systemin immune network within *Nicotiana*.

## Results

### *Nicotiana* contains PS genes encoding SYS peptides

While PS genes within other members of the *Solanoideae* have been known for over two decades (Constabel et al. 1998), no known PS ortholog outside of *Solanoideae* has been previously described. To find and annotate PS orthologs within *Nicotiana*, we used the phylogenomic analysis pipeline BLAST-Align-Tree (BAT) (Sheppard et al. 2026). Although many hits shared little to no similarity with known SlPS sequences, we found that SlPS1 formed an outgroup to a clade containing SlPS2-5 (Fig. 1A). An unpruned version of the BAT tree can be found in Fig. S1. Three *N. tabacum* genes and one *N. benthamiana* gene cluster with SlPS2-5 and include 2 features of other PS genes: the 27-aa N-terminal motif of SlPS2/3/4 and a conserved C-terminal peptide (Fig. 1A, Fig. S2). Conserved sequence similarity was highest to SlPS4, so we named them NtPS4A/B/C and NbPS4. Notably, the gene encoding SlPro-antiSYS is not annotated in ITAG4.0 (Wang et al. 2025) and was not included in this analysis.

**Figure 1:**
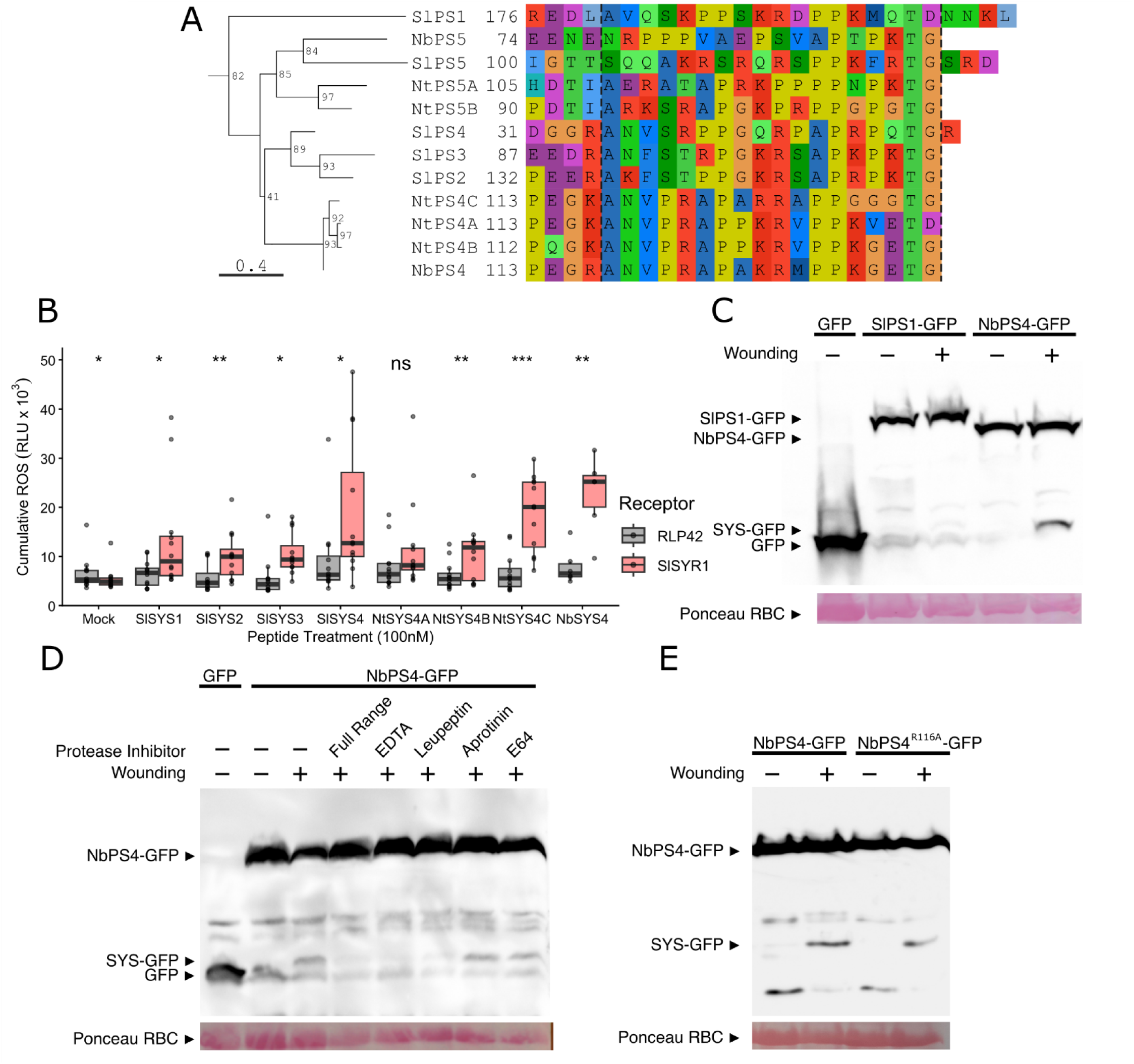
Nicotiana contains PS homologs which encode bioactive SYS peptides, including NbPS4 which is processed in response to wounding via a mechanism distinct from SlPS1 processing. (A) Phylogenetic tree of PS sequences from *S. lycopersicum*, *N. benthamiana* and *N. tabacum* and an alignment of their C-termini. The tree was built using the BAT phylogenomics pipeline (Sheppard et al. 2026) using SlPS4 and SlPS1 CDS sequences as BLAST queries against reference genomes. PS names correspond to gene IDs as follows: SlPS1: Solyc05g051750.3.1, SlPS2: Solyc04g011735.1.1, SlPS3: Solyc04g011750.3.1, SlPS4: Solyc04g011760.3.1, SlPS5: Solyc04g011770.2.1, NtPS4A: NC_134102.1_cds_XP_016473095.1_83812, NtPS4B: NC_134093.1_cds_XP_016458871.1_52941, NtPS4C: NC_134095.1_cds_XP_016439249.2_59210, NtPS5A:NC_134095.1_cds_XP_016439247.2_59212, NtPS5B: NC_134095.1_cds_XP_016439246.1_59213, NbPS4: Nbe.v1.1.chr04g32890, NbPS5: Nbe.v1.1.chr14g18230. Numbers indicate the position of the first displayed AA. Dashed lines indicate either the known (SlPS1), or putative cut sites. (B) Cumulative ROS of *N. benthamiana* expressing either RLP42 or SlSYR1 under a 35S promoter after being treated with 100 nM flg22, SlSYS1-4, NtSYS4A-C, or NbSYS4. The cumulative ROS was calculated by summing every minute’s ROS measurement from minute 5 to 40. Points indicate the average of technical replicates for each biological replicate (n=12). Statistical significance was calculated via paired students t test between RLP42 and SlSYR1 samples of the same biological replicates with significance denoted by * = p<0.05, ** = p<0.01, and *** = p<0.001. (C, D and E) Western blots of N. benthamiana tissue expressing either GFP, SlPS1-GFP, NbPS4-GFP or NbPS4^R116A^ -GFP. Samples indicated as wounded were incubated at 25 °C for 5 minutes before protein extraction. Protease inhibitors were added via infiltration before samples were taken. All blots used the same anti-GFP primary antibody, and a Ponceau S stain of the Rubisco band was included as a loading control. Similar results for each blot were seen in at least two independent experiments.

To determine whether these new *Nicotiana* systemins were bioactive, we synthesized their putative SYS peptides (Fig. 1A, Table S1) and tested them alongside SlSYS1-4 against SlSYR1. We found that all tested SYS peptides except NtSYS4A elicited ROS production in a SlSYR1-dependent manner, when compared to an unrelated control receptor, RLP42 (Fig. 1B).

To test whether *Nicotiana* PS proteins are processed in the same wounding dependent manner as SlPS1-4, we expressed SlPS1-GFP and NbPS4-GFP under a 35S promoter in *N. benthamiana*. We then wounded the agroinfiltrated area by incubating crushed tissue at 25 °C for 5 minutes before extracting protein. Full length NbPS4-GFP accumulated in both wounded and unwounded samples, and wounding induced a smaller band consistent with the size of NbSYS4-GFP, indicating a cleavage event near the C-terminus (Fig. 1C). While SlPS1 accumulated well, it did not process in response to wounding, indicating that cleavage of SlPS1 in tomato and NbPS4 in *N. benthamiana* likely occurs via distinct mechanisms (Fig. 1C).

Cleavage of NbPS4-GFP is attenuated in samples containing a broad-spectrum protease inhibitor, indicating that NbPS4 processing is protease dependent. In addition, the calcium chelator EDTA and the serine, cysteine, and threonine protease inhibitor leupeptin also attenuated processing, but not the serine protease inhibitor aprotinin, or the cysteine protease inhibitor E64 (Fig. 1D).

SlPS1 processing was previously shown to be phytaspase-mediated, and dependent on aspartate residues at positions 179 and 196 (Beloshistov et al. 2018). However, the SlPS2, SlPS4 and *Nicotiana* PS sequences lack these key residues. We noticed that there is a conserved arginine/lysine residue at position 116, just before the putative cut site. The processing of the phytocytokine AtPROPEP1 was previously shown to be dependent on an arginine residue on the N-terminal side of the cut site (Hander et al. 2019), leading us to hypothesize that this residue might be implicated in the processing of NbPS4. However, we found that NbPS4^R116A^-GFP processed comparably well to NbPS4-GFP (Fig. 1E), indicating that another mechanism of processing is sufficient.

### LAB *N. benthamiana* lacks a functional SYR and cannot detect NbSYS4

Publicly available data provide evidence for *NbPS4* transcription, with highest expression in the roots (Fig. S3A), leading us to suspect that NbPS4 may play an active role in root defense signaling. However, we found that NbSYS4 did not elicit a ROS response in either *N. benthamiana* leaves (Fig. 2A) or roots (Fig. 2B). These negative results led us to more closely investigate SYRs within *Nicotiana*.

**Figure 2.**
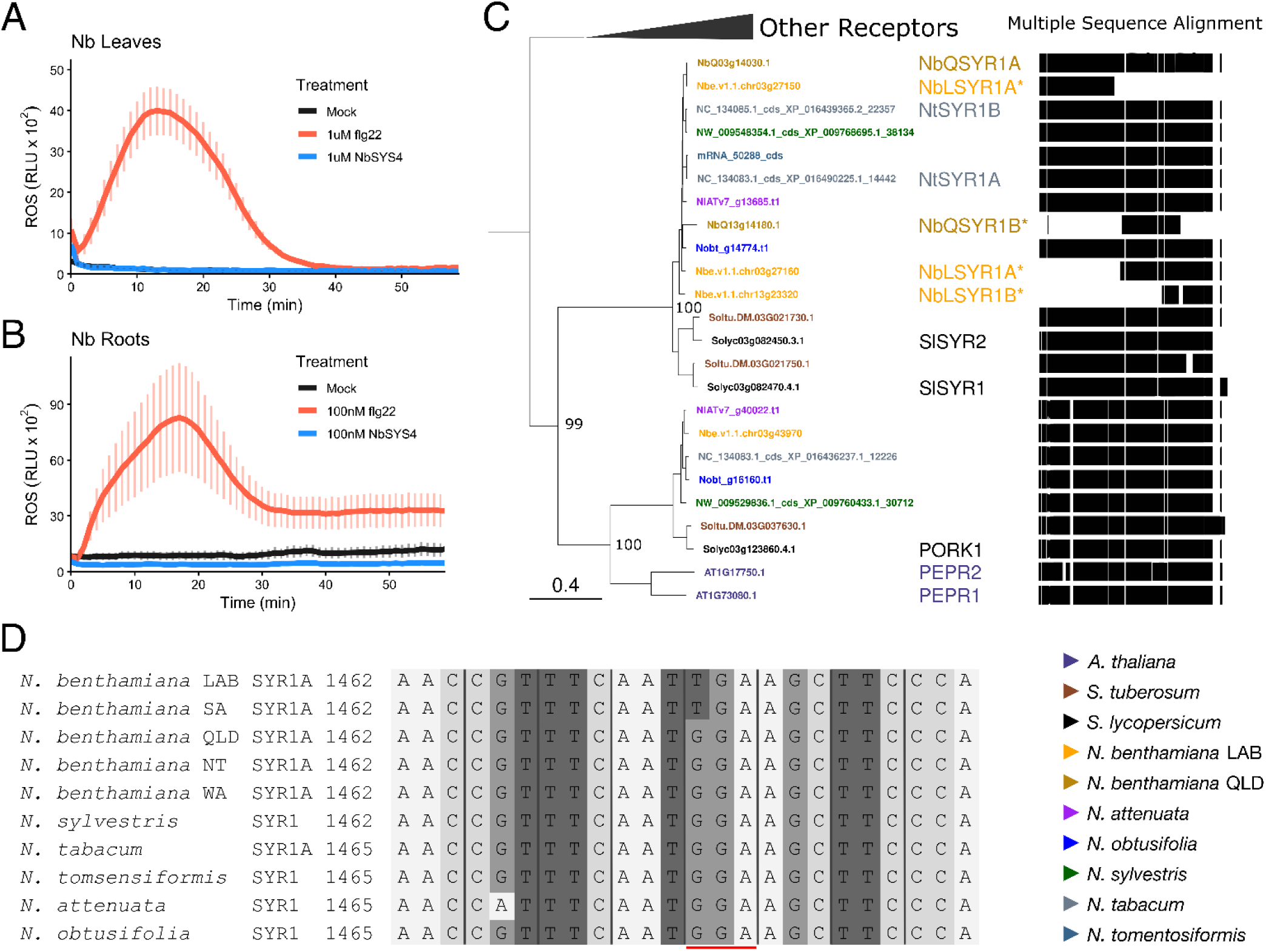
LAB *N. benthamiana* cannot detect NbSYS4 and contains a premature stop codon in NbSYR1A. (A and B) ROS time course of *N. benthamiana* leaves against 1 µM flg22 and NbSYS4 (A) and roots against 100 nM flg22 and NbSYS4 (B). Neither *N. benthamiana* leaves nor roots exhibit a ROS burst in response to NbSYS4. Each line consists of the average from all technical replicates across all biological replicates (A: n=6, B: n=36). Error bars are the SEM of the average of the technical replicates for each biological replicate. (C) Phylogenetic tree of SYRs and related receptors. Tree was built using the BAT phylogenomics pipeline using NbQSYR1A CDS (NbQ03g14030.1) as the query sequence against reference genomes of the 8 species and 2 accessions in the legend. Black bars in the Multiple Sequence Alignment (MSA) indicate the presence of aa residues after alignment; gaps in the MSA indicate gaps in the aa alignment. An asterisk next to a gene name indicates a suspected pseudogene based on MSA. Bootstraps are shown as values from 0 to 100. (D) A subset of a CDS alignment of SYRs from 5 *N. benthamiana* accessions and 5 *Nicotiana* species. Black lines indicate the reading frame and the lower red line indicates the location of a SNP found in the LAB and SA *N. benthamiana* accessions which introduces a premature stop codon. The numbers before each sequence indicate the location of the start of each displayed sequence in the larger CDS.

We used the BAT phylogenomics pipeline to find SlSYR1 homologs in *A. thaliana*, *S. tuberosum, S. lycopersicum*, and 6 *Nicotiana* species covering the breadth of the genus, including both the widely used LAB and wild Queensland (QLD) accessions of *N. benthamiana*. As in previous studies (Wang et al. 2018), the SYRs formed a high confidence sister clade to PEPRs (Fig. 2C, Fig. 4S). As expected (Wang et al. 2018), *Solanum* species contained 2 SYR genes, the diploid *Nicotiana* species contained 1 full length SYR, and the allotetraploid *N. tabacum* contained two. Both *N. benthamiana LAB (Nbe_v1.1)* and *N. benthamiana* QLD (NbQld183.v103) contained one truncated SYR we named NbSYR1B. *N. benthamiana LAB* contained two additional SYR associated annotations,, Nbe.v1.1.chr03g27150 and Nbe.v1.1.chr03g27160 which are adjacent on the chromosome but encode separate complementary parts of a single SYR (Fig. 2C, Fig. 5S). Aligning the combined gene model (NbLSYR1A) with the SYR1s from the other *Nicotiana* species’ SYR1As, as well as the other accessions from within *N. benthamiana* revealed a mutation causing a premature stop codon in the LRR ectodomain, which was found in LAB and SA accessions of *N. benthamiana* (Fig. 2D). Notably, SA was previously shown to be the wild population most closely related to LAB and also contains the same nonfunctional allele of the RDR1 gene as LAB (Bally et al. 2015). Phylogenomic analysis thus indicates a novel stop codon in *SYR1A* in LAB and SA accessions of *N. benthamiana*.

### *N. tabacum roots* are responsive to NbSYS4

While *N. benthamiana* was unresponsive to NbSYS4, the presence of multiple full length SYR1 genes in *N. tabacum* (Fig. 2C) implied that it may be responsive to NbSYS4. Based on transcriptional data (Fig. S3A), we tested both mature *N. tabacum* leaves and seedling roots with NbSYS4 via a ROS assay. Although NtSYR1A has a similar expression pattern to SlSYR1 (Fig. S3B), NbSYS4 failed to induce a ROS response in *N. tabacum* leaves (Fig. 3A). However, we did see a ROS response in *N. tabacum* roots, consistent with the presence of a functional SYR in *N. tabacum* (Fig. 3B) To further validate this *N. tabacum* phenotype we treated *N. tabacum and N. benthamiana* seedling roots with NbSYS4, SlSYS4 and flg22. In *N. benthamiana,* neither SYS peptide had any measurable impact on root growth while flg22 had a strong inhibitory effect (Fig. 3C and 3D). *N. tabacum* seedlings were insensitive to both SlSYS4 and flg22, but responsive to NbSYS4 (Fig. 3E and 3F). The insensitivity to flg22 was highly unexpected given that it induced a ROS response in *N. tabacum* roots (Fig. 3B), but may be due to degradation, dosage, or sensitivity issues over the seven day length of the assay. We conclude that *N. tabacum* seedlings respond to NbSYS4, consistent with the presence of intact SYR1 homologs in the tobacco genome.

**Figure 3.**
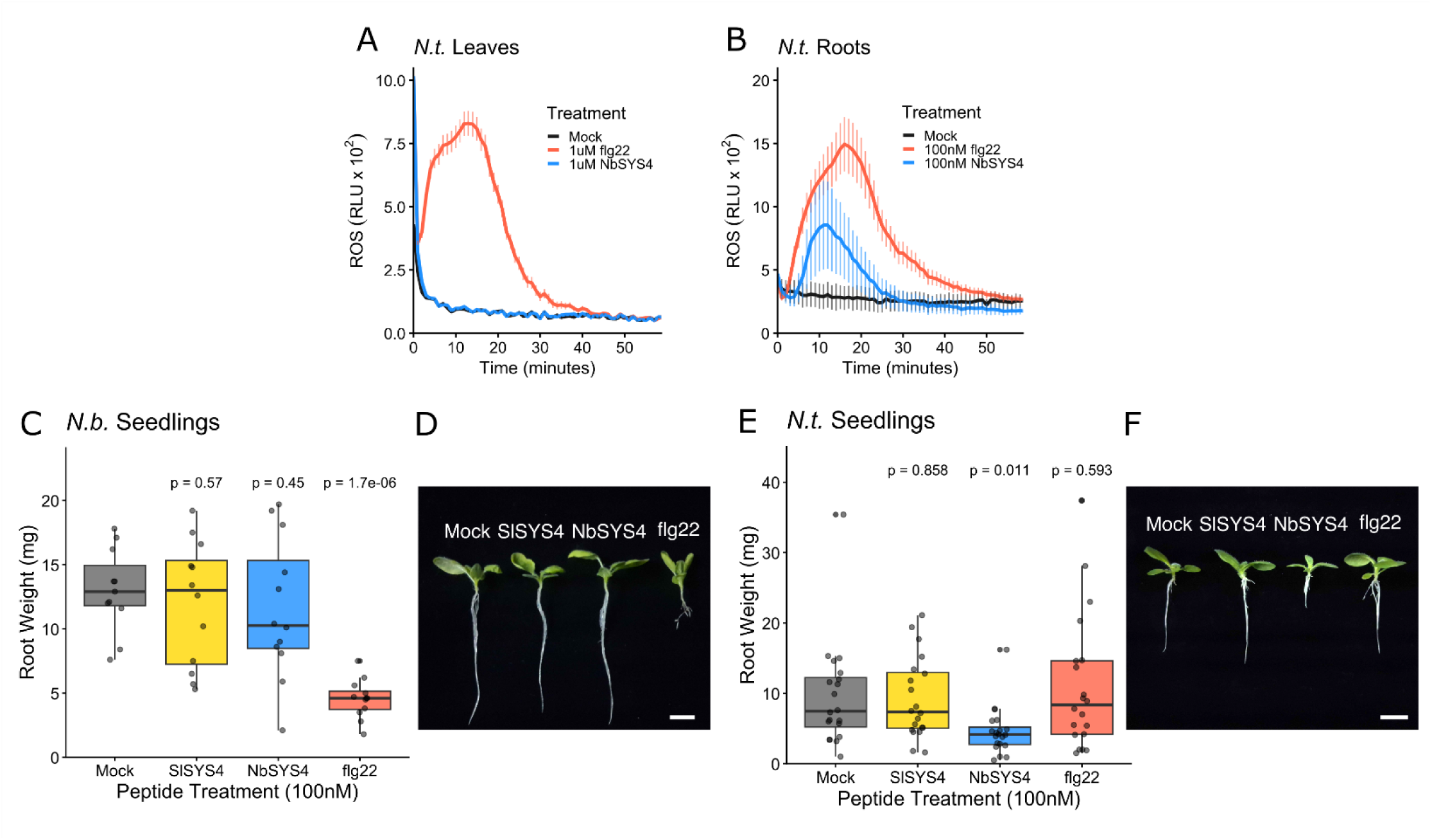
*N. tabacum* roots exhibit multiple immune phenotypes in response to NbSYS4. (A and B) ROS time course of *N. tabacum* leaves against 1uM flg22 and NbSYS4 (A) and roots against 100nM flg22 and NbSYS4 (B). Each line consists of the average from all technical replicates across all biological replicates (A: n=4, B: n=48). Error bars are the SEM of the average of the technical replicates for each biological replicate. (C and E) The root weight of *N. benthamiana* (C) or *N. tabacum* (E) after being grown in 1x liquid MS for 7 days with the indicated peptide treatment. 12-20 plants were measured per treatment. (D and F) Photos taken of representative seedlings taken from C and E respectively. Scale bar represents 1 cm. Error bars in A and B are SEM. P-values in C and E were calculated via unpaired Students T-test between the given peptide treatment and the mock.

### Expressing NtSYR1A in *N. benthamiana* leaves confers the ability to detect SYS peptides

We investigated if the putative NtSYR1s were involved in NbSYS4 detection, by characterizing the more highly expressed NtSYR1A. We tested if the transient expression of NtSYR1A in *N. benthamiana* could confer the ability to detect SYS peptides. We validated NtSYR1A accumulation by tagging it with GFP and probing for it on a Western Blot (Fig. S6). Expressing NtSYR1A in LAB *N. benthamiana* enabled NbSYS4 and SlSYS4 induced ROS responses (Fig. 4A). Other SYS peptides (SlSYS1-3 and NtSYS4A-C) also induced ROS responses against NtSYR1A expressing *N. benthamiana* to varying degrees (Fig. 4B). Notably the NtSYS4A ROS response was weaker than the other SYS elicitors, mirroring its behavior against SlSYR1 (Fig. 4B).

**Figure 4.**
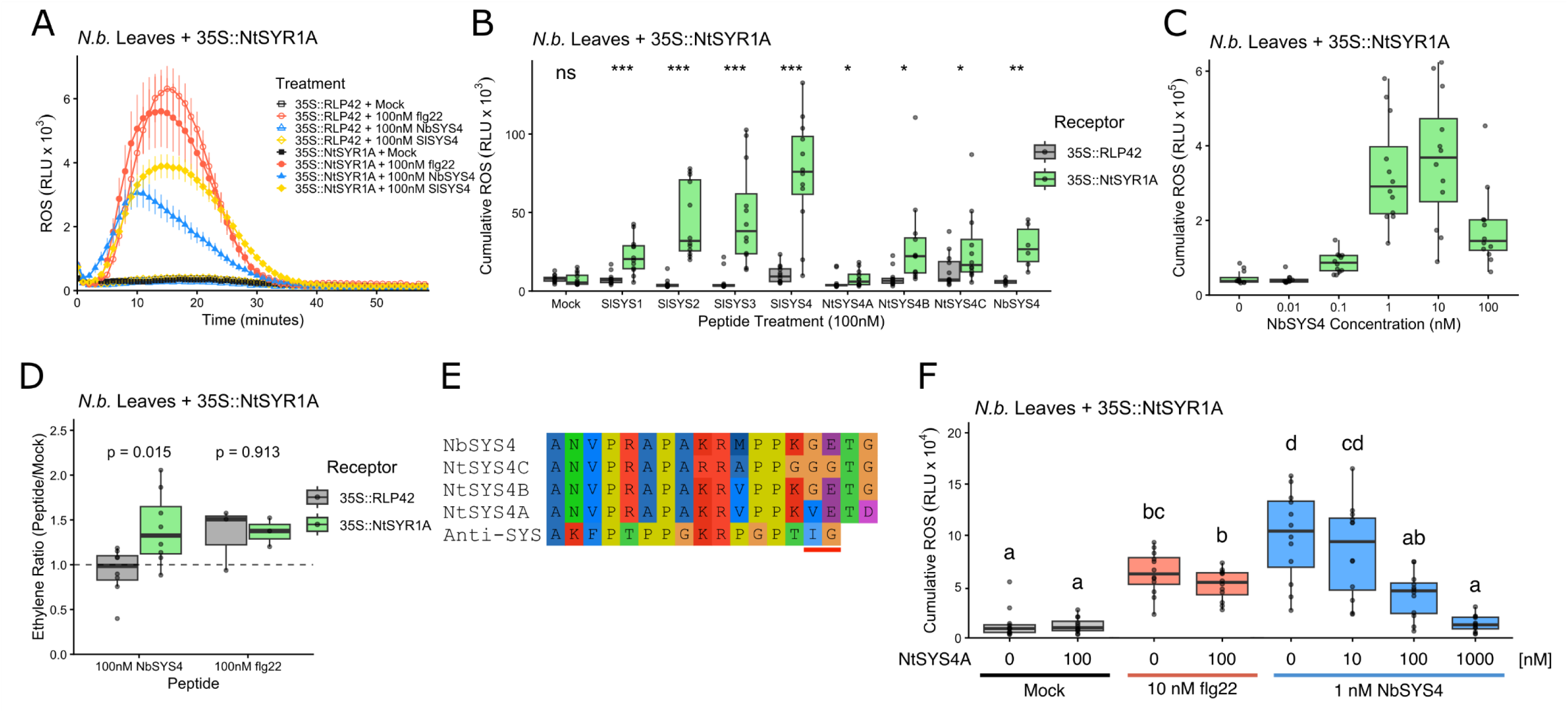
Expressing NtSYR1A in *N. benthamiana* enables varied immune responses to different SYS peptides. (A) ROS time course of *N. benthamiana* expressing either RLP42 (negative control), or NtSYR1A after being treated with a Mock solution, 100 nM flg22, 100 nM NbSYS4 and 100nM SlSYS4. Each line consists of an average of all technical replicates from all biological replicates (n = 8). (B) Cumulative ROS of *N. benthamiana* expressing either RLP42 or NtSYR1A after being treated with a Mock solution or 100nM of the indicated peptide. Points indicate the average of technical replicates for each biological replicate (n=12). Statistical significance was calculated via paired Student’s t test between RLP42 and NtSYR1A samples of the same biological replicates with significance denoted by * = p<0.05, ** = p<0.01, and *** = p<0.001. (C) Cumulative ROS of *N. benthamiana* expressing NtSYR1A when treated with the indicated concentration of NbSYS4. Points indicate the average of technical replicates for each biological replicate (n=12). (D) Ratio of ethylene produced between the mock and indicated peptide treatment in RLP42 and NtSYR1A expressing *N. benthamiana*. Dashed line indicates a peptide treatment to mock ratio of 1. Points indicate the average of technical replicates for each biological replicate (NbSYS4: n=11, flg22: n=3). (E) Alignment of *Nicotiana* systemins and SlAntiSYS. Red bar indicates residues key in deciding normal SYS vs. antiSYS activity. (F) Cumulative ROS of NtSYR1A expressing *N. benthamiana* after being pretreated 15 min before by the indicated concentration of NtSYS4A then treated with either flg22 or NbSYS4 at the indicated concentration. Points indicate the average of technical replicates for each biological replicate (n=12). (B, C, and F) The cumulative ROS was found by summing every minute’s RLU measurement from minute 5 to 40. Error bars in A are SEM. Significance in B and the p values in D were calculated by paired student’s T-test between the RLP42 and NtSYR1A samples of the same biological replicate for each peptide with significance denoted by * = p<0.05, ** = p<0.01, and *** = p<0.001. Statistically distinct groups in F were determined by ANOVA followed by Tukey’s HSD and are indicated by letters..

To see if the sensitivity of NtSYR1A was comparable to that previously demonstrated by SlSYR1 (Wang et al. 2018) we tested various concentrations of NbSYS4 against NtSYR1A expressing LAB *N. benthamiana*. Like SlSYS1 against SlSYR1, NbSYS4 induced a ROS response at sub-nanomolar concentrations, reaching a near maximal response at around 1 nM, and decreasing at 100 nM, indicating that NtSYR1A is likely a high affinity receptor (Fig. 4C). In addition, NtSYR1A but not RLP42 enabled an NbSYS4 induced ethylene response (Fig. 4D). Since 100 nM NtSYS4A caused only a weak response against NtSYR1A and an insignificant response against SlSYR1, we further analyzed its amino acid sequence. NtSYS4A carries V15 and D18, where NtSYS4B carries G15 and G18 (Fig. 4E). It has been previously shown that SlSYS1 M15I and Q16G substitutions (amino acids from SlAntiSYS) are sufficient to cause an inhibitory phenotype (Wang et al. 2025). Similar C-terminal differences in NtSYS4A are consistent with potential antiSYS activity. Pretreating LAB *N. benthamiana* expressing NtSYR1A with NtSYS4A causes a dose-dependent inhibition of the ROS response to NbSYS4, but not flg22, in a manner consistent with the inhibition of SlSYS1 by SlAntiSYS (Wang et al. 2025; Fig. 4F). Thus, sequence differences among NtSYS genes may provide mechanisms for regulation.

## Discussion

The systemin family of phytocytokines has been shown to be a key regulator of the immune response within *Solanoideae*, regulating responses to a wide range of pests and pathogens (Skripnikov et al. 2026, Wang et al. 2025, Ryan et al. 1992). Here we identified and described multiple systemins and their receptors within *Nicotiana*, the first instance of systemins being described outside of *Solanoideae.* Specifically, we highlighted NbPS4, NtPS4A, NtPS4B, and NtPS4C, all of which contain peptides that share substantial sequence similarities with SlSYSes and are recognized by SlSYR1. We also showed that NbPS4 is processed in a similar manner to SlSYSes. In our investigation of *Nicotiana* SYRs we found that NbLSYR1A, the SlSYR1 ortholog in LAB accession *N. benthamiana*, contains premature stop codons, providing a potential explanation as to why neither SlSYSes nor any of the *Nicotiana* SYSes tested caused any response in LAB *N. benthamiana*. This lineage specific loss of SYR1 was supported by the related *N. tabacum* both responding to NbSYS4, and containing NtSYR1A, which, when expressed in *N. benthamiana*, conferred the ability to detect systemins.

We also uncovered multiple functional differences within and between SlSYS and *Nicotiana* SYS peptides. We found that SlSYS4 causes a strong ROS response against NtSYR1A in the *N. benthamiana* expression system but surprisingly lacks activity against *N. tabacum* root growth. This is potentially due to our synthetic peptide stock of SlSYS4 containing an additional C-terminal arginine, which has been shown to be cleaved during processing in tomato (Wang et al. 2026) but may not be processed in intact *N. tabacum* roots. We also showed that NtSYS4A causes a very weak ROS response compared to other SYSes, and that pretreating leaves with it causes an attenuation of an NbSYS4 ROS response in a dose dependent manner, a phenotype similar to that of SlAntiSYS (Wang et al. 2025). NtPS4A also has a much higher level of constitutive transcription than NtPS4B or NtPS4C, a pattern also held by SlPro-antiSYS and SlPS2/4. Our findings suggest that there is substantial species-specificity to regulation of immune responses by different systemin peptide repertoires.

Interestingly, most SYRs, including SlSYR1, appear to be expressed in roots far more strongly than in leaves (Fig. S3B). SlPS2/3/4 have previously been shown to be primarily root expressed (Wang et al. 2025), and NtPS4A, the only NtPS with detectable expression, is also primarily root expressed. *N. tabacum* also only exhibited responses to NbSYS4 in root tissue (Fig. 3B and 3E). Together, this suggests that the ancestral state of SYS signaling may be a root-based signal, and that SlPS1 being primarily a leaf-based signal is indicative of a more derived state.

Interestingly, both NbPS4 and NbLSYR1A are still transcribed at moderate levels within LAB *N. benthamiana* (Fig. S3A and S3B). While this may simply be a transcriptional remnant of a recent mutation, there is potential for a non-canonical function of NbPS4 and the pseudogene NbLSYR1A.

We identified a premature stop codon in the NbSYR1A of LAB and SA *N. benthamiana* accessions, but intact CDS sequences in other varieties and *Nicotiana* species. LAB *N. benthamiana* first rose to prominence as a model organism in part due to its extremely poor immune system and fast life cycle (Goodin et al. 2008). Previous studies have found function impairing mutations in *RDR1*, a viral defense gene (Bally et al. 2015) and *Chr10-LDOX*, a gene involved in anthocyanin production (Ahmed et al. 2026), which are found in LAB and SA *N. benthamiana* but are absent from the majority of wild accessions. Holding with the growth defense trade-off hypothesis (Huot et al. 2014), LAB and SA have substantially faster life cycles than other populations (Bally et al. 2015). The lack of a functional SYR in these populations can be viewed in this framework of prioritizing vigor at the expense of defense. Similarly, SYR1 was previously shown to negatively contribute to yield when compared to *syr1* knockout plants (Zhou et al. 2025). That there are three separate instances of defense related genes being rendered non-functional within the LAB/SA populations indicate the likely existence of more waiting to be uncovered.

Our results provide additional context to the widely held assumption that LAB *N. benthamiana* cannot detect systemins. Rather than *Nicotiana* being a clade lacking systemins, our findings suggest that LAB *N. benthamiana* specifically has lost the ability to detect systemins. As a model organism that appears to be SYR1 deficient, but closely related to SYR1 containing accessions, LAB *N. benthamiana* is likely to have retained the necessary co-receptors and downstream signaling partners necessary for biologically relevant SYR functions, making it an ideal *tabula rasa* for further investigations into systemin signaling.

The conserved SYR1 receptor and PS homologs in *Nicotiana* indicate that SYR-systemin signaling is conserved in *Solanaceae*, and that both SYRs and PS have undergone gene family expansions in *Solanum* and *Nicotiana*. Outside of *Solanaceae*, SYR1-like receptors can be found as far back as monocots (Wang et al. 2018), but we found no obvious PS orthologs. This indicates that the evolutionary origins of the systemin SYR system are far from understood, and that additional studies into non-solanaceous SYS-SYR1 systems are needed.

Further work on the systemin and SYR families is needed. While this study has introduced several *Nicotiana* PS, and found surface level similarities in response to *Solanoideae* PS genes, whether they have similar downstream transcriptional and hormonal impacts is yet to be uncovered. Additionally, signaling differences between *Nicotiana* SYS peptides, and how conserved those responses are across wild diploid species needs further work. Previous studies have shown that the N-terminal propeptide sequence of SlPS1 has bioactive components (Molisso et al. 2022, Criscuolo et al. 2025) but the roles that the propeptide sequence plays in all other PS proteins, both in localization and processing, and as a bioactive component itself, are yet to be investigated.

## Methods

### Plant growth

All plants were grown in Conviron PGC Flex growth chambers set to 12-h light/12-h dark, 60% relative humidity, 25 °C, and a light intensity of 350 µmol m^-2^ s^-1^.

### Cloning

Leaf, root and stem tissue of *N. benthamiana* and *N. tabacum* was harvested and ground using liquid N_2_ chilled balls in a 30 Hz shaker. RNA was extracted using Trizol (Invitrogen) and cDNA libraries were created using the SMARTer® RACE 5’/3’ Kit (Takara Bio). NtSYR1A-GFP was constructed by PCR amplifying (Q5 NEB hot start) the NtSYR1A sequence from cDNA then cloning it into the Gateway vector pGM, then transferring to pEarleyGate 103 (Early et al. 2006). RLP42-GFP was previously constructed in the same manner. All other constructs were created via Golden Gate hierarchical modular cloning (Weber et al. 2011, Engler et al. 2014). The NbPS4 sequence was PCR amplified from cDNA, while SlSYR1 and SlPS1 were amplified from a domesticated TWIST fragment. Due to an error in SlSYR1 domestication it was amplified in 3 fragments. The resulting PCR fragments were ligated into the L0 vector pACH9121. The NbPS4^R116A^ was created by amplifying the L0 NbPS4 construct using Q5 PCR with the mutation introduced in the primers, then treating the resulting product with the KLD enzyme mixture (NEB). The L0 constructs were then ligated into an L1 construct using a modified pGreenII (Hellens et al. 2000) backbone including BsaI cutsites, 35S Cauliflower Mosaic Virus +5’UTR Tobacco mosaic virus (pICH51266), C-terminal GFP (Aequorea victoria; pICSL50008), and the OCS1 terminator (pICH41432). Construct accuracy was checked at the L0 stage via Sanger sequencing and final expression plasmids were whole plasmid sequenced via Plasmidsaurus. All primers used can be found in Table S2.

### Peptides

All peptides used in this study were synthesized by GenScript at a crude purity, and reconstituted in H_2_O . Precise peptide sequences for flg22, SlSYS1-4, NtSYS4A/B/C and NbSYS4 can be found in Table S1.

### *N. benthamiana* transient expression

Constructs were electroporated into *Agrobacterium tumefaciens* strain GV3101. Overnight cultures were resuspended in 150 μM acetosyringone in 10 mM 2-(N-morpholino) ethanesulfonic acid (MES), pH 5.6, and 10 mM MgCl_2_. After 3 hr of incubation at room temperature, *N. benthamiana* leaves of 4 to 5 week-old plants were infiltrated with an optical density at 600 nm (OD_600_) of 1 for PS-GFP constructs, or 0.3 for SYR/RLP42-GFP constructs.

### ROS assay

Leaf punches were taken with a 4 mm biopsy punch. Seedling roots were taken from 10-14 day old seedlings sprouted on 1x MS, and cut just below the hypocotyl. Both were then floated in 150 μL of H2O using individual cells of a white 96-well white bottom plate(BRANDplates F pureGrade S white [REF 781665]) with the leaf punches being adaxial side up. After overnight incubation in the same growth conditions as the original plants, with the seedling root plates being covered, ROS production was measured immediately after the addition of a 100 μL assay solution which contained 10 μg/mL horseradish peroxidase, 10 mM L-012 and the peptide treatment at 2.5x the final indicated concentration. Luminescence was quantified with a TECAN SPARK plate reader every minute for an hour using an integration time of 500 ms. Leaf-based experiments used 4-12 distinct bioreps, with experiments testing multiple receptors including both receptors in distinct zones on the same leaf. For each bioreplicate 3-6 individual technical replicates were designated for use with each peptide treatment, so the pre-ROS treatment plates were functionally identical within a given experiment. For seedling root ROS each individual seedling root was placed in each well. For the NtSYS4A inhibition assay, 50 μL of 4x the final indicated concentration of NtSYS4A was added 15 minutes prior to the addition of 50 μL of the assay solution, which contained 20 μg/mL luminol-horseradish peroxidase, 20 mM L-012 and the peptide treatment at 5x the final indicated concentration.

### Western blots

48 hours post agro-infiltration, samples were taken with a 17 mm punch then immediately frozen in liquid N_2_. Samples being treated with a protease inhibitor or being directly compared to them were infiltrated immediately prior to punching with either a mock H2O solution, or the corresponding protease inhibitor at the following concentrations, Pierce Complete Protease Inhibitor (1X), EDTA (50 mM), leupeptin (20 μM), aprotinin (1 μM), E64 (100 μM). Samples were then cryogenically ground into a fine powder. Samples that were wounded were then incubated at 25 °C for 5 minutes, then immediately refrozen in liquid N_2_. Next, 120 μL of 90 °C 3X LL buffer was added to each sample, they were vortexed for 1 minute, then incubated at 90 °C for 10 minutes. Samples were centrifuged at > 12,000 g and the supernatant was collected, and separated by size on a 12.5% SDS-PAGE gel, then wet transferred overnight to a nitrocellulose membrane. The membrane was then blocked using 5% non-fat dry milk in 1X TBST (20 mM Tris, 150 mM NaCl, 0.1% Tween) for 1 hour, then washed once in 1X TBST, then incubated with mouse anti-GFP (1:1,000; Roche: 11814460001) in 1X TBST for 1 hour, then washed with 1X TBST twice, then incubated for 1 hour with anti-mouse IgG:HRP (1:10,000; AB_228307) in 1X TBST, and washed a final time with 1X TBST. Membranes were then imaged using a combination of SuperSignal™ West Pico PLUS Chemiluminescent Substrate (Thermo Fisher) and SuperSignal™ West Femto PLUS Chemiluminescent Substrate (Thermo Fisher) in a 9:1 ratio. After imaging (via AZI600-01), Ponceau stain was used to ensure roughly equal protein loading.

### Data analysis and figure creation

All data analysis and graphing were performed in R (4.5.1) using RStudio (2025.05.1+513). The package tidyverse (2.0.0) was used for data manipulation, and the packages ggplot2 (4.0.2) and ggpubr (0.6.3) were used for graphing and statistical analysis. Full figures were created in Inkscape (1.4.4).

### BAT tree construction

The tool BLAST-Align-Tree (Sheppard et al. 2026) was used to create phylogenetic trees and alignments of both PS and SYR sequences. The genomes used for both the PS and SYR trees can be found in Table S3. The PS tree was created by querying cds annotations using the CDS sequence of SlPS1 (Solyc05g051750.3.1), SlPS2 (Solyc04g011735.1.1), and NbPS4 (Nbe.v1.1.chr04g32890) and pulling 12 sequences from each genome. The sequences were aligned using MAFFT (v7.427; Katoh & Standley 2013) in L-INS-i mode with otherwise default settings. The tree was imported into iTOL (v7.6; Letunic & Bork 2024) where nodes that obviously lacked sequence homology were manually pruned; full unedited trees can be found in the figure S1. The SYR1/PEPR tree was created by querying cds annotations against the NbQSYR1A CDS (NbQ03g14030.1) pulling 20 sequences per genome, with the aligner set to MAFFT in L-INS-i mode with otherwise default settings.

### Transcriptional data

Transcriptional data for the PS and SYR1 genes found in figure S3. were sourced from Ding (2025) for *N. tabacum,* the Tomato Functional Genomics Database (TFGD), experiment D004 IDs S0592 (Heinz root) and S0593 (Heinz leaves) for *S. lycopersicum* (Tomato Genome Consortium 2012), and from the leaf and root tracks on https://apollo.nbenth.com/annotator/index (Ranakawa et al. 2023) for LAB and QLD *N. benthamiana*. Average root and leaf expression for *N. tabacum* genes was found by taking the mean of the TPM for that given gene across all samples of that tissue type. To normalize the Fragments per Kilobase (FPKM) of the TFGD data to TPM, we used the following formula TPM = Mean Transcript Length * RPKM * 10^6^ / Sum of RPKM. To calculate TPM for the LAB and QLD *N. benthamiana* using the raw counts we found the counts for our gene of interest then TPM was calculated using the following formula TPM = (Counts/(Total Reads/10^6^))*(Average transcript length/transcript of interest length).

### Root Growth Inhibition Assay

*N. benthamiana* and *N. tabacum* seedlings were sown on 1x MS agar plates 8 and 13 days prior to the assay respectively and kept in the same growth chamber as the other *N. benthamiana*. An ⅛” drill bit was used to drill 4 holes in the lids of 5 mL Falcon tubes. The tubes were then filled to the brim with 1x MS containing 100 nM of either flg22, SlSYS4, NbSYS4 or a mock treatment. The seedlings were then carefully transferred to the tubes, with each tube containing 4 seedlings. The tubes were placed under a humidity dome and returned to the growth chamber where the plates were incubated. After 7 days the seedlings were removed, the roots were removed at the hypocotyl via a razor blade, then patted dry with a Kimwipe and weighed on a balance.

### Ethylene Assay

Entire, fully expanded 5 week old *N. benthamiana* leaves were agroinfiltrated with either NtSYR1A-GFP or RLP42-GFP. At 48 hpi, half of each leaf was infiltrated with H_2_O then immediately dried and four 10-mm punches were taken from the infiltrated half of the leaf and placed into a 5 mL tube which was sealed with a rubber cap. This was then repeated on the other half of the leaf using either 100 nM flg22 or 100 nM NbSYS4. Three hours after the tubes were sealed 1 mL was pulled from the headspace. Ethylene content was measured using a gas chromatography machine (HP 5890 series 2, Supelco #13,018 U, and 80/100 Hayesep Q 3 ft × ⅛ in. x 2.1 mm nickel). The Treatment/Mock ratio was calculated by finding the ratio of the peak height of the peptide treated sample to the mock treated sample for each individual leaf.

## Acknowledgements

This research was funded by the NIH 5R35GM151272 grant to A.D.S

## Author contributions

Conceptualization: A.D.S., E.W.M.; Investigation A.D.S., E.W.M.; Visualization: E.W.M., Writing - Original Draft: E.W.M.; Writing - Review & Editing: A.D.S, E.W.M.; Funding acquisition: A.D.S.

---

**Figure S1:**
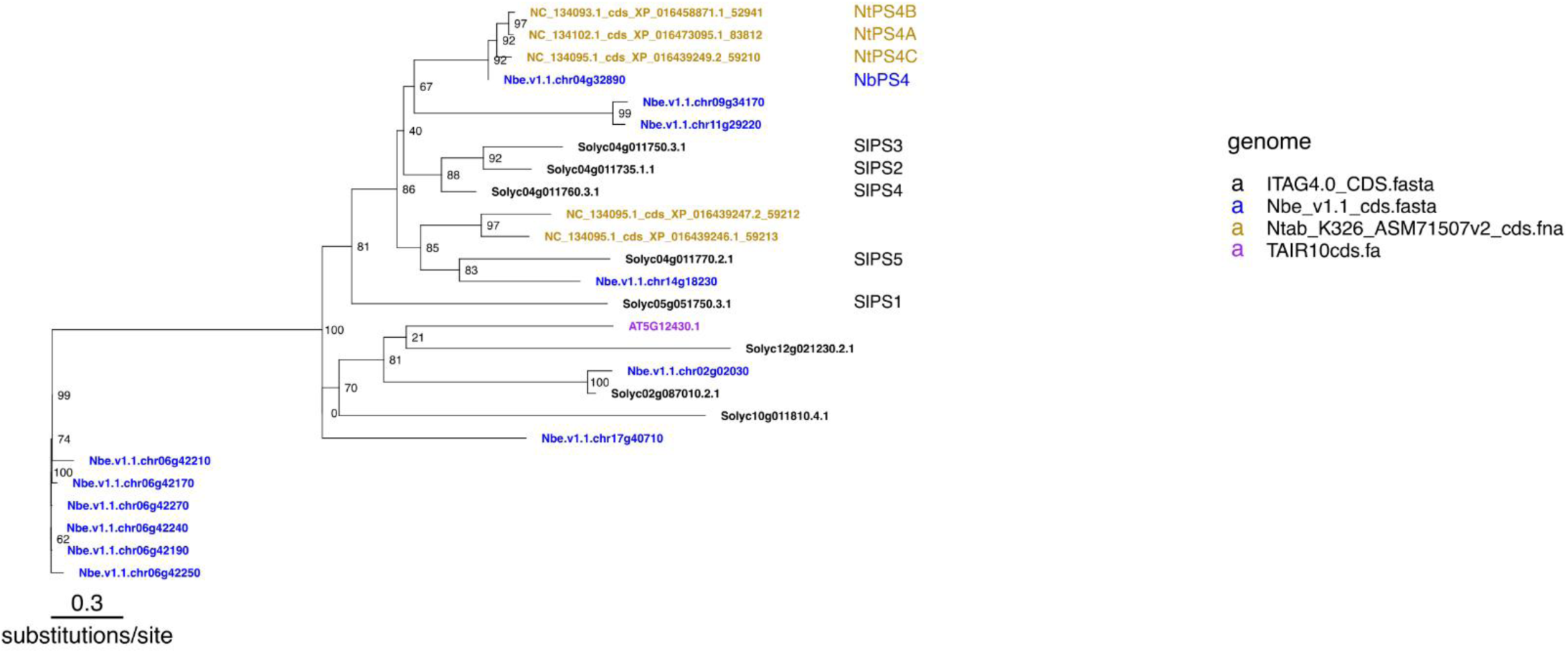
Unedited BAT tree of PS genes in *S. lycopersicum*, *N. tabacum*, and *N. benthamiana*. Numbers in the phylogeny indicate bootstrap values. BAT was performed using the genomes listed on the right, using SlPS1(Solyc05g051750) and SlPS2 (Solyc04g011735) as query sequences, pulling 10 queries from Nbe_v1.1_cds.fasta and 5 from other genomes. The sequence aligner was set to MAFFT on mode L-INS-i.

**Figure S2:**
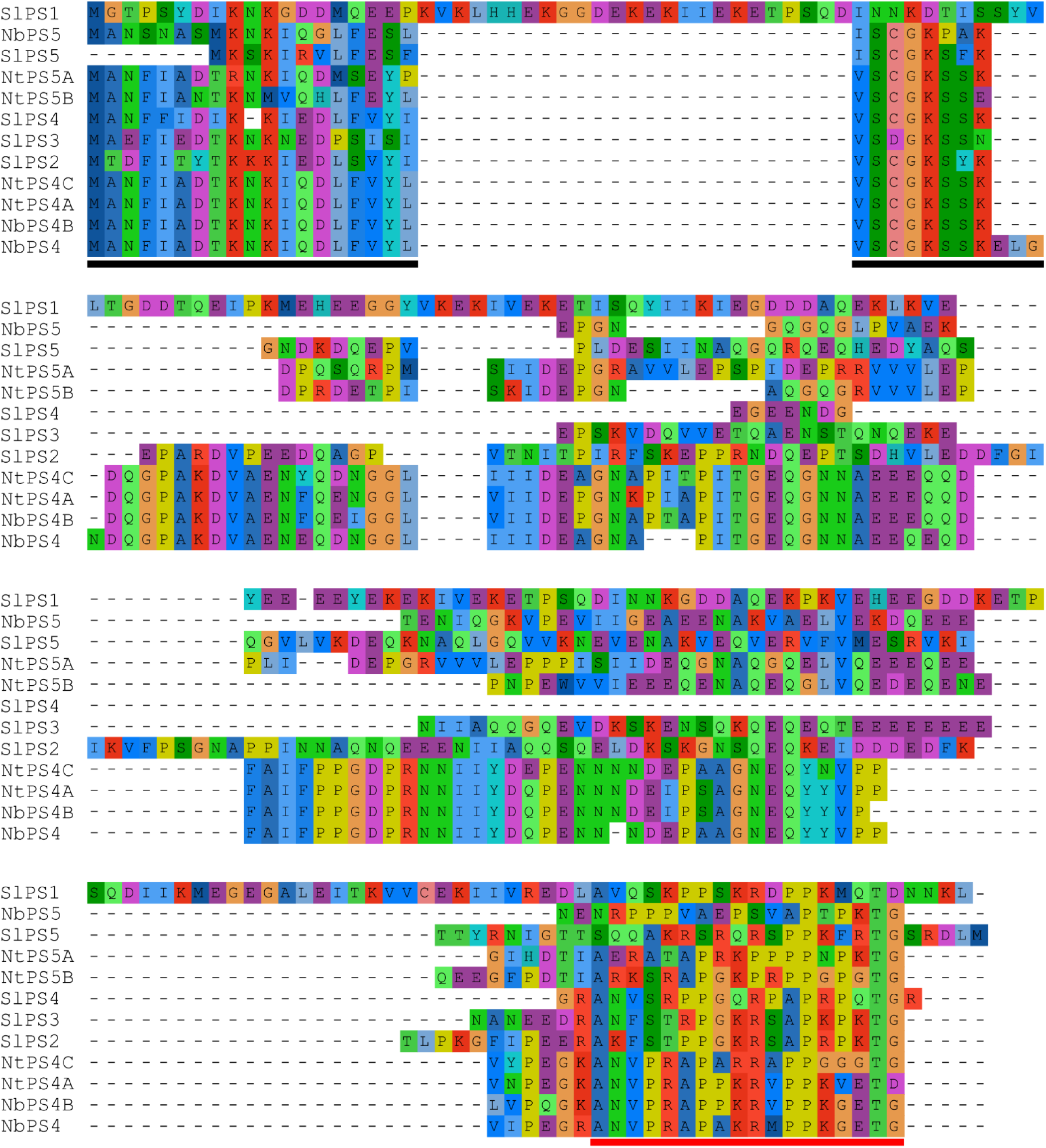
Full aa sequence alignment of PS proteins in figure 1A. Black underline highlights a conserved 27 amino acid N-terminal domain found in all SlPS2/3/4, NtPS4A/B/C and NbPS4. The conserved C-terminal peptide is underlined in red. Sequences were visualized and aligned in Aliview using Muscle 3.8 with minor manual tweaks. PS names correspond to gene IDs as follows: SlPS1: Solyc05g051750, SlPS2: Solyc04g011735.1.1, SlPS3: Solyc04g011750.3.1, SlPS4: Solyc04g011760.3.1, SlPS5: Solyc04g011770.2.1, NtPS4A: NC_134102.1_cds_XP_016473095.1_83812, NtPS4B: NC_134093.1_cds_XP_016458871.1_52941, NtPS4C: NC_134095.1_cds_XP_016439249.2_59210, NtPS5A:NC_134095.1_cds_XP_016439247.2_59212, NtPS5B: NC_134095.1_cds_XP_016439246.1_59213, NbPS4: Nbe.v1.1.chr04g32890, NbPS5: Nbe.v1.1.chr14g18230.

**Figure S3:**
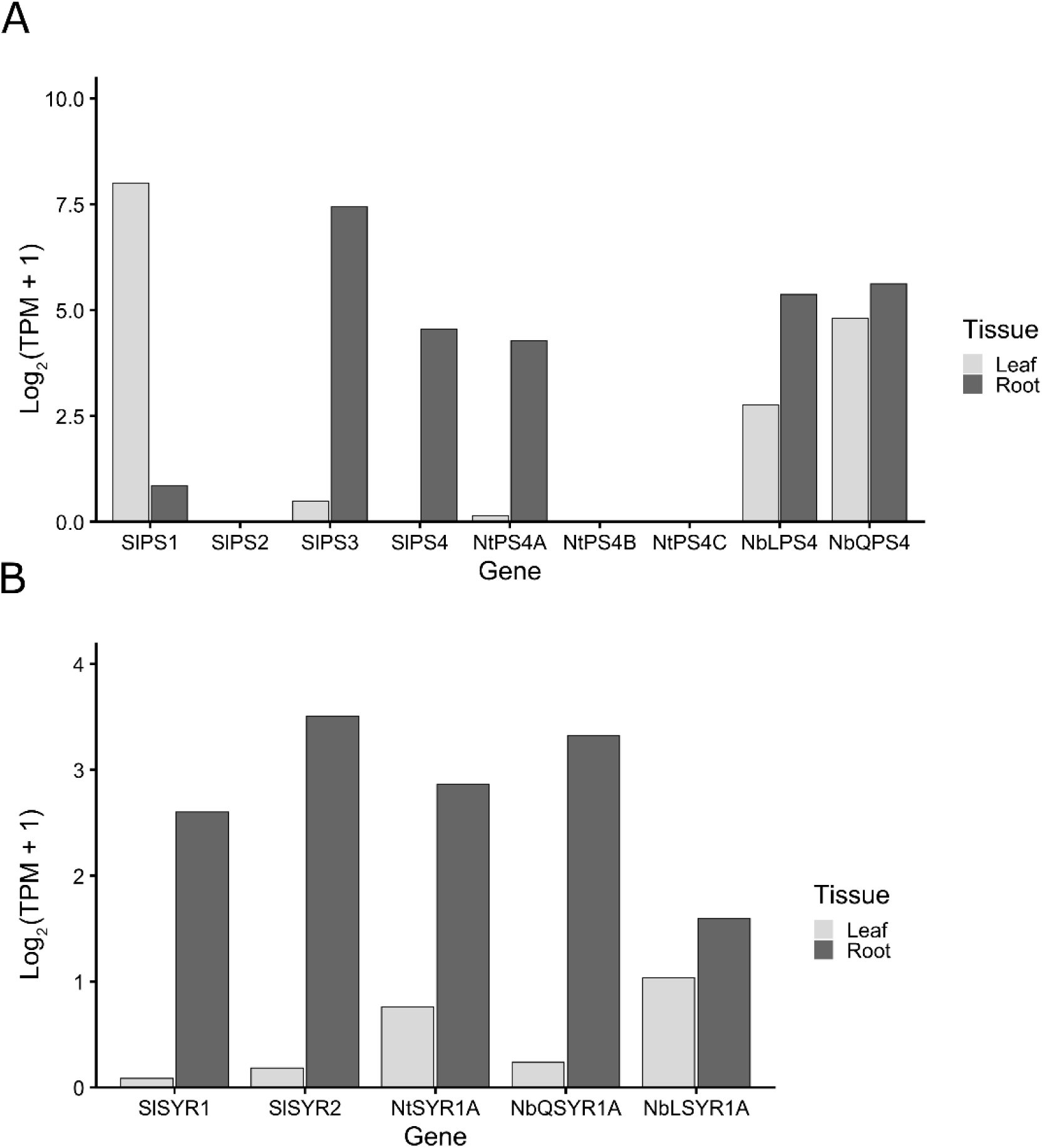
Transcriptional levels of PS and SYR genes in *S. lycopersicum*, *N. tabacum*, and *N. benthamiana*. (A) Expression of various PS genes in mature leaves and roots. Shown as a log2 transformation of Transcript per Kilobase Million (TPM). (B) Expression of various SYR1s in mature leaves and roots. Shown as a log2 transformation of Transcript per Kilobase Million (TPM).

**Figure S4:**
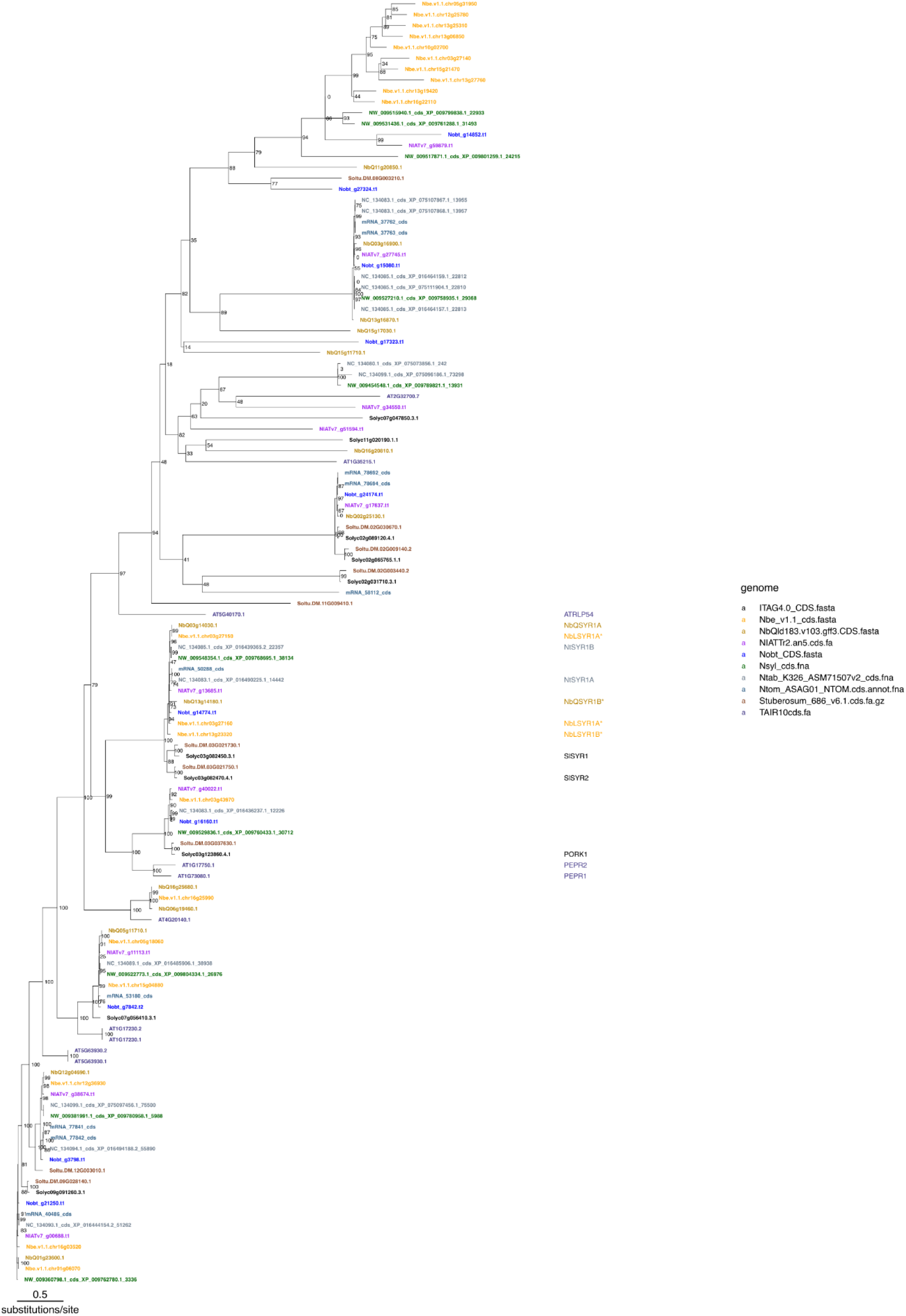
Unedited BAT tree of SYR family. Numbers in the phylogeny indicate bootstrap values. BAT was performed using the genomes listed on the right, using NbQSYR1 (NbQ03g14030.1) as the main query sequence, pulling 20 queries from Nbe_v1.1_cds.fasta and 10 from other genomes. The sequence aligner was set to MAFFT on mode L-INS-i.

**Figure S5:**
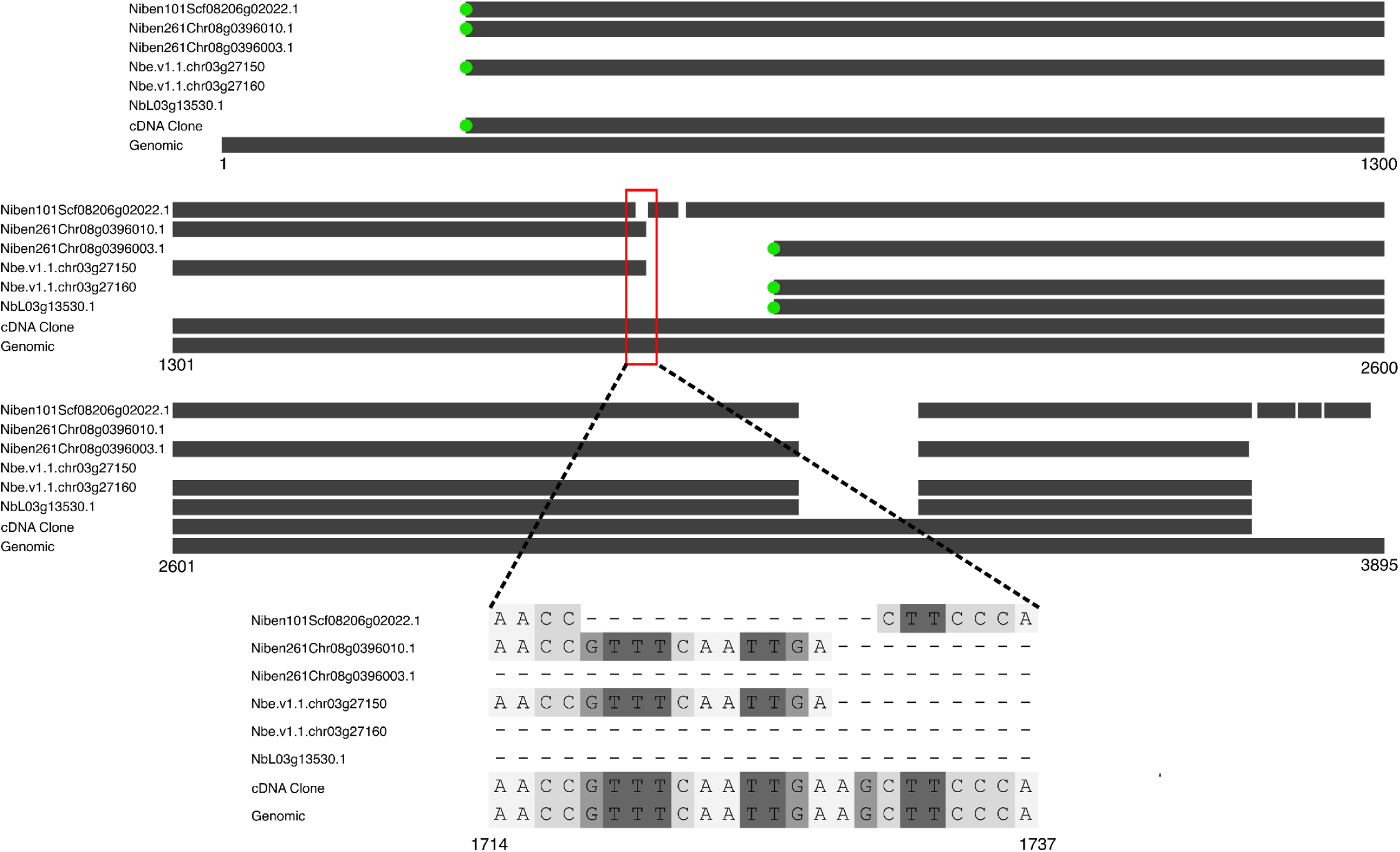
Alignment of CDS annotations from different *N. benthamiana* genomes to cloned cDNA and genomic sequences. Sequence alignment of annotations of NbSYR1A from Niben v1.0.1, Niben v2.6.1, Nbe.v1.1 and Lab360. Green dots indicate the start of the CDS annotation. Also included in the alignment is the sequencing results from our attempt to clone NbSYR1A from cDNA (primers in Table S2), and the genomic sequence corresponding to the unspliced mRNA transcript, taken from Nbe.v1.1 and validated against Lab360. The red box indicates the portion of the sequence shown in detail in figure 2D and below. Numbers indicate the nucleotide position of genomic sequence.

**Figure S6:**
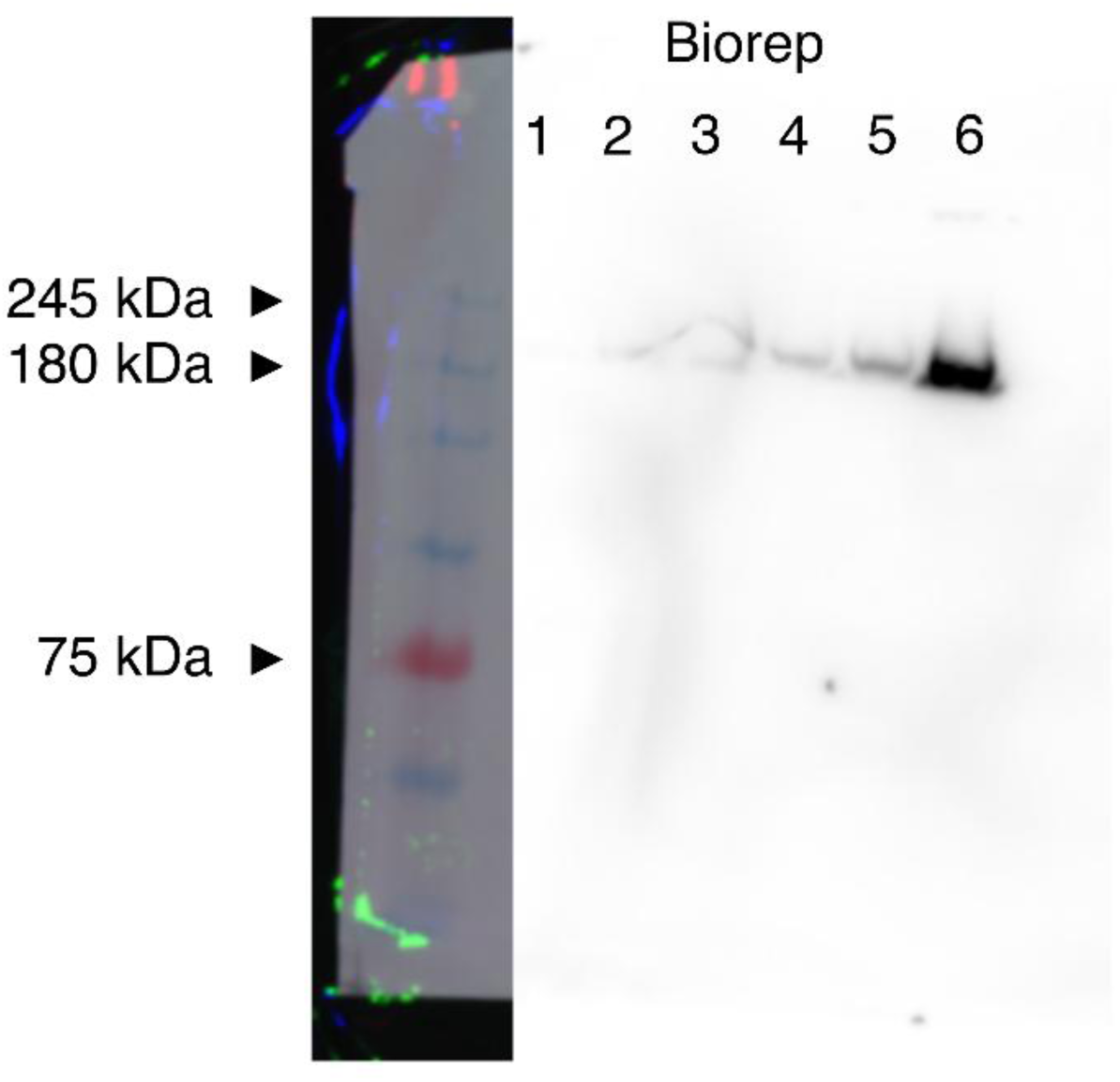
Western blot of 35S::NtSYR1A-GFP expressing *N. benthamiana*. Tissue taken from leaf regions used for ROS assays in figure 4A and 4B. Each lane represents combined tissue from two leaves on the same plant. anti-GFP primary antibody was used. Based on previous expression of SlSYR1-GFP in *N. benthamiana* (Zhou et al. 2025), a band of ∼180 kDa was expected for NtSYR1A-GFP. The Ladder shown is Prometheus ProSignal® Full-Range Prestained Protein Ladder.

## References

1. Ahmed, Z., An, J., Hayashi, S., Bally, J., Winefield, C., & Waterhouse, P. M. (2026). Harnessing Bulk-Segregant Mapping to Identify Trait-Associated Genes in the Allopolyploid Model Plant Nicotiana benthamiana. Plant Biotechnology Journal, 24(11), 5575–5587. 10.1111/pbi.70560

2. Bally, J., Jung, H., Mortimer, C., Naim, F., Philips, J. G., Hellens, R., Bombarely, A., Goodin, M. M., & Waterhouse, P. M. (2018). The Rise and Rise of Nicotiana benthamiana: A Plant for All Reasons. Annual Review of Phytopathology, 56, 405–426. 10.1146/annurev-phyto-080417-050141

3. Bally, J., Nakasugi, K., Jia, F., Jung, H., Ho, S. Y. W., Wong, M., Paul, C. M., Naim, F., Wood, C. C., Crowhurst, R. N., Hellens, R. P., Dale, J. L., & Waterhouse, P. M. (2015). The extremophile Nicotiana benthamiana has traded viral defence for early vigour. Nature Plants, 1(11), 15165. 10.1038/nplants.2015.165

4. Beloshistov, R. E., Dreizler, K., Galiullina, R. A., Tuzhikov, A. I., Serebryakova, M. V., Reichardt, S., Shaw, J., Taliansky, M. E., Pfannstiel, J., Chichkova, N. V., Stintzi, A., Schaller, A., & Vartapetian, A. B. (2018). Phytaspase-mediated precursor processing and maturation of the wound hormone systemin. New Phytologist, 218(3), 1167–1178. 10.1111/nph.14568

5. Buonanno, M., Coppola, M., Di Lelio, I., Molisso, D., Leone, M., Pennacchio, F., Langella, E., Rao, R., & Monti, S. M. (2018). Prosystemin, a prohormone that modulates plant defense barriers, is an intrinsically disordered protein. Protein Science, 27(3), 620–632. 10.1002/pro.3348

6. Chakraborty, A., & Xu, S. (2026). Chromosome-level genome assemblies of Nicotiana attenuata (coyote tobacco) and Nicotiana obtusifolia (desert tobacco). Scientific Data, 13(1), 441. 10.1038/s41597-026-07080-y

7. Constabel, C. P., Yip, L., & Ryan, C. A. (1998). Prosystemin from potato, black nightshade, and bell pepper: Primary structure and biological activity of predicted systemin polypeptides. Plant Molecular Biology, 36(1), 55–62. 10.1023/A:1005986004615

8. Criscuolo, M. C., Magliulo, R., Castaldi, V., Cirillo, V., Cristiani, C., Negroni, A., Aprile, A. M., Molisso, D., Buonanno, M., Esposito, D., Langella, E., Monti, S. M., & Rao, R. (2025). Salt Stress Mitigation and Field-Relevant Biostimulant Activity of Prosystemin Protein Fragments: Novel Tools for Cutting-Edge Solutions in Agriculture. Plants, 14(15), 2411. 10.3390/plants14152411

9. Ding, Y. (2025). *Gene expression level in different samples* [Data set]. figshare. 10.6084/m9.figshare.29666060.v1

10. Earley, K. W., Haag, J. R., Pontes, O., Opper, K., Juehne, T., Song, K., & Pikaard, C. S. (2006). Gateway-compatible vectors for plant functional genomics and proteomics. The Plant Journal, 45(4), 616–629. 10.1111/j.1365-313X.2005.02617.x

11. Engler, C., Youles, M., Gruetzner, R., Ehnert, T.-M., Werner, S., Jones, J. D. G., Patron, N. J., & Marillonnet, S. (2014). A Golden Gate Modular Cloning Toolbox for Plants. ACS Synthetic Biology, 3(11), 839–843. 10.1021/sb4001504

12. Goodin, M. M., Zaitlin, D., Naidu, R. A., & Lommel, S. A. (2008). Nicotiana benthamiana: Its History and Future as a Model for Plant–Pathogen Interactions. Molecular Plant-Microbe Interactions, 21(8), 1015–1026. 10.1094/MPMI-21-8-1015

13. Hander, T., Fernández-Fernández, Á. D., Kumpf, R. P., Willems, P., Schatowitz, H., Rombaut, D., Staes, A., Nolf, J., Pottie, R., Yao, P., Gonçalves, A., Pavie, B., Boller, T., Gevaert, K., Van Breusegem, F., Bartels, S., & Stael, S. (2019). Damage on plants activates Ca2+-dependent metacaspases for release of immunomodulatory peptides. Science, 363(6433), eaar7486. 10.1126/science.aar7486

14. Hellens, R. P., Edwards, E. A., Leyland, N. R., Bean, S., & Mullineaux, P. M. (2000). pGreen: A versatile and flexible binary Ti vector for Agrobacterium-mediated plant transformation. Plant Molecular Biology, 42(6), 819–832. 10.1023/A:1006496308160

15. Hou, S., Liu, D., & He, P. (2021). Phytocytokines function as immunological modulators of plant immunity. Stress Biology, 1(1), 8. 10.1007/s44154-021-00009-y

16. Huffaker, A., Pearce, G., Veyrat, N., Erb, M., Turlings, T. C. J., Sartor, R., Shen, Z., Briggs, S. P., Vaughan, M. M., Alborn, H. T., Teal, P. E. A., & Schmelz, E. A. (2013). Plant elicitor peptides are conserved signals regulating direct and indirect antiherbivore defense. Proceedings of the National Academy of Sciences, 110(14), 5707–5712. 10.1073/pnas.1214668110

17. Huot, B., Yao, J., Montgomery, B. L., & He, S. Y. (2014). Growth–Defense Tradeoffs in Plants: A Balancing Act to Optimize Fitness. Molecular Plant, 7(8), 1267–1287. 10.1093/mp/ssu049

18. Katoh, K., & Standley, D. M. (2013). MAFFT Multiple Sequence Alignment Software Version 7: Improvements in Performance and Usability. Molecular Biology and Evolution, 30(4), 772–780. 10.1093/molbev/mst010

19. Kurotani, K., Hirakawa, H., Shirasawa, K., Tagiri, K., Mori, M., Ramadan, A., Ichihashi, Y., Suzuki, T., Tanizawa, Y., An, J., Winefield, C., Waterhouse, P. M., Miura, K., Nakamura, Y., Isobe, S., & Notaguchi, M. (2025). Establishing a comprehensive web-based analysis platform for Nicotiana benthamiana genome and transcriptome. The Plant Journal, 121(1), e17178. 10.1111/tpj.17178

20. Lamesch, P., Berardini, T. Z., Li, D., Swarbreck, D., Wilks, C., Sasidharan, R., Muller, R., Dreher, K., Alexander, D. L., Garcia-Hernandez, M., Karthikeyan, A. S., Lee, C. H., Nelson, W. D., Ploetz, L., Singh, S., Wensel, A., & Huala, E. (2012). The Arabidopsis Information Resource (TAIR): Improved gene annotation and new tools. Nucleic Acids Research, 40(D1), D1202–D1210. 10.1093/nar/gkr1090

21. Letunic, I., & Bork, P. (2024). Interactive Tree of Life (iTOL) v6: Recent updates to the phylogenetic tree display and annotation tool. Nucleic Acids Research, 52(W1), W78–W82. 10.1093/nar/gkae268

22. Liu, L., Liu, X., Bai, Z., Tanveer, M., Zhang, Y., Chen, W., Shabala, S., & Huang, L. (2024). Small but powerful: RALF peptides in plant adaptive and developmental responses. Plant Science, 343, 112085. 10.1016/j.plantsci.2024.112085

23. Molisso, D., Coppola, M., Buonanno, M., Di Lelio, I., Aprile, A. M., Langella, E., Rigano, M. M., Francesca, S., Chiaiese, P., Palmieri, G., Tatè, R., Sinno, M., Barra, E., Becchimanzi, A., Monti, S. M., Pennacchio, F., & Rao, R. (2022). Not Only Systemin: Prosystemin Harbors Other Active Regions Able to Protect Tomato Plants. Frontiers in Plant Science, 13. 10.3389/fpls.2022.887674

24. Pearce, G., Strydom, D., Johnson, S., & Ryan, C. A. (1991). A Polypeptide from Tomato Leaves Induces Wound-Inducible Proteinase Inhibitor Proteins. Science. 10.1126/science.253.5022.895

25. Pham, G. M., Hamilton, J. P., Wood, J. C., Burke, J. T., Zhao, H., Vaillancourt, B., Ou, S., Jiang, J., & Buell, C. R. (2020). Construction of a chromosome-scale long-read reference genome assembly for potato. GigaScience, 9(9), giaa100. 10.1093/gigascience/giaa100

26. Ranawaka, B., An, J., Lorenc, M. T., Jung, H., Sulli, M., Aprea, G., Roden, S., Llaca, V., Hayashi, S., Asadyar, L., LeBlanc, Z., Ahmed, Z., Naim, F., de Campos, S. B., Cooper, T., de Felippes, F. F., Dong, P., Zhong, S., Garcia-Carpintero, V., … Waterhouse, P. M. (2023). A multi-omic Nicotiana benthamiana resource for fundamental research and biotechnology. Nature Plants, 9(9), 1558–1571. 10.1038/s41477-023-01489-8

27. Rhodes, J., Yang, H., Moussu, S., Boutrot, F., Santiago, J., & Zipfel, C. (2021). Perception of a divergent family of phytocytokines by the Arabidopsis receptor kinase MIK2. Nature Communications, 12(1), 705. 10.1038/s41467-021-20932-y

28. Schiavinato, M., Marcet-Houben, M., Dohm, J. C., Gabaldón, T., & Himmelbauer, H. (2020). Parental origin of the allotetraploid tobacco Nicotiana benthamiana. The Plant Journal, 102(3), 541–554. 10.1111/tpj.14648

29. Sheppard, B. D., Behnken, B., & Steinbrenner, A. (2026). BAT: An integrated pipeline for gene tree construction, annotation, and functional inference (p. 2026.05.07.721474). bioRxiv. 10.64898/2026.05.07.721474

30. Sierro, N., Auberson, M., Dulize, R., & Ivanov, N. V. (2024). Chromosome-level genome assemblies of Nicotiana tabacum, Nicotiana sylvestris, and Nicotiana tomentosiformis. Scientific Data, 11(1), 135. 10.1038/s41597-024-02965-2

31. Sierro, N., Battey, J. N. D., Ouadi, S., Bakaher, N., Bovet, L., Willig, A., Goepfert, S., Peitsch, M. C., & Ivanov, N. V. (2014). The tobacco genome sequence and its comparison with those of tomato and potato. Nature Communications, 5(1), 3833. 10.1038/ncomms4833

32. Sierro, N., Battey, J. N., Ouadi, S., Bovet, L., Goepfert, S., Bakaher, N., Peitsch, M. C., & Ivanov, N. V. (2013). Reference genomes and transcriptomes of Nicotiana sylvestris and Nicotiana tomentosiformis. Genome Biology, 14(6), R60. 10.1186/gb-2013-14-6-r60

33. Skripnikov, A., Suprunova, T., Taliansky, M., & Kalinina, N. O. (2026). Synthetic Potato Systemin Enhances Solanum tuberosum Resistance to Potato Virus Y. Biomolecules, 16(9), 1353. 10.3390/biom16091353

34. The Tomato Genome Consortium. (2012). The tomato genome sequence provides insights into fleshy fruit evolution. Nature, 485(7400), 635–641. 10.1038/nature11119

35. Wang, L., Einig, E., Almeida-Trapp, M., Albert, M., Fliegmann, J., Mithöfer, A., Kalbacher, H., & Felix, G. (2018). The systemin receptor SYR1 enhances resistance of tomato against herbivorous insects. Nature Plants, 4(3), 152–156. 10.1038/s41477-018-0106-0

36. Wang, L., Maier, L.-P., Pham, N., Wang, Y. L., Wang, X., Schaller, A., Fliegmann, J., Erb, M., Boller, T., & Felix, G. (2025). A receptor antagonist counterbalances multiple systemin phytocytokines in tomato. Cell. 10.1016/j.cell.2025.07.044

37. Wang, X., Yuan, W., Pham, N.-T., Wang, L., Li, R., Steppuhn, A., Weigel, D., Stintzi, A., Felix, G., & Schaller, A. (2026). Recruitment of SERK co-receptors determines signaling specificity within the systemin peptide family (p. 2026.02.13.705698). bioRxiv. 10.64898/2026.02.13.705698

38. Weber, E., Engler, C., Gruetzner, R., Werner, S., & Marillonnet, S. (2011). A Modular Cloning System for Standardized Assembly of Multigene Constructs. PLOS ONE, 6(2), e16765. 10.1371/journal.pone.0016765

39. Xu, S., Brockmöller, T., Navarro-Quezada, A., Kuhl, H., Gase, K., Ling, Z., Zhou, W., Kreitzer, C., Stanke, M., Tang, H., Lyons, E., Pandey, P., Pandey, S. P., Timmermann, B., Gaquerel, E., & Baldwin, I. T. (2017). Wild tobacco genomes reveal the evolution of nicotine biosynthesis. Proceedings of the National Academy of Sciences, 114(23), 6133–6138. 10.1073/pnas.1700073114

40. Yamaguchi, Y., Barona, G., Ryan, C. A., & Pearce, G. (2011). GmPep914, an Eight-Amino Acid Peptide Isolated from Soybean Leaves, Activates Defense-Related Genes. Plant Physiology, 156(2), 932–942. 10.1104/pp.111.173096

41. Zhou, K., Wu, F., Deng, L., Xiao, Y., Yang, W., Zhao, J., Wang, Q., Chang, Z., Zhai, H., Sun, C., Han, H., Du, M., Chen, Q., Yan, J., Xin, P., Chu, J., Han, Z., Chai, J., Howe, G. A., … Li, C. (2025). Antagonistic systemin receptors integrate the activation and attenuation of systemic wound signaling in tomato. Developmental Cell, 60(4), 535–550.e8. 10.1016/j.devcel.2024.11.005

42. Ziemann, S., van der Linde, K., Lahrmann, U., Acar, B., Kaschani, F., Colby, T., Kaiser, M., Ding, Y., Schmelz, E., Huffaker, A., Holton, N., Zipfel, C., & Doehlemann, G. (2018). An apoplastic peptide activates salicylic acid signalling in maize. Nature Plants, 4(3), 172–180. 10.1038/s41477-018-0116-y

